# Individual vocal recognition in the black-headed spider monkey (*Ateles fusciceps*)

**DOI:** 10.1101/2023.04.19.537610

**Authors:** D. Nagle, T. Power, J. L. Quinn, C. A. Troisi

**Author notes:** Corresponding author: Camille Troisi –.

## Abstract

Individual vocal recognition – the ability to discriminate between individuals based on sound – is thought to be particularly useful for social species that regularly lose visual contact between group members. For instance, individuals living in a fission-fusion group that feed on patchily distributed food in a complex physical environment (e.g., dense forest) are likely to depend heavily on vocalisation to identify individuals that are good at finding food. Spider monkeys (*Ateles spp*.) live in such environments and have distinctive individual contact calls (*whinnies*). We used a habituation-dishabituation playback-paradigm to investigate whether black-headed spider monkeys (*A. fusciceps*) are able to discriminate between female individuals in their group. We found that a group of captive spider monkeys was able to discriminate between individuals using those contact calls. Although many primate species have been found to have individual characteristics in their calls, this is one of the few direct experimental evidence of vocal recognition using a habituation-dishabituation paradigm.

## Introduction

One crucial aspect of any social interaction is the possibility of recognising and distinguishing specific individuals such as mates, offspring, allies or competitors (Steiger & Müller, 2008; Tibbetts et al., 2008; Tibbetts & Dale, 2007). Recognising others can be based on individual-level or class-level characteristics (Tibbetts & Dale, 2007). Class-level recognition occurs when individuals identify others based on class characteristics, such as age, sex, or familiarity, while individual-level recognition occurs when identification is based on distinctive individual characteristics (Dale et al., 2001). In this process, an individual’s characteristics are associated with unique information about the identity of the signalling individual, which allows to later recognize the individual following a learning process over repeated encounters (Tibbetts & Dale, 2007). Some individuals can also recognise specific individuals from other species (Johnson et al., 2018). Individuals signal their identity based on various modalities such as chemical, visual and acoustic signals, and numerous species from diverse taxa have been shown to use various forms of individual recognition.

One of the best known examples of individual recognition using sounds is the ability of emperor penguins to relocate their chicks among thousands using vocal recognition (Aubin et al., 2000). Individual recognition by means of vocalisation is critically important when individuals are separated by visual contact from their group and allows for effective communication between these individuals (Santorelli et al., 2013; Spillmann et al., 2010). Complex visual environments (e.g. dense vegetation, patchy distribution of resource) or fission-fusion societies place high demands on mechanisms that allow individuals to coordinate movement, identify subgroups, and keep in contact with group members while foraging (Milton, 2000). In particular, individual recognition based on vocal cues are well known to strengthen group cohesion or facilitate social decisions in many species (Kondo & Watanabe, 2009).

Using vocal cues, primates are able to distinguish between familiar and unfamiliar individuals (e.g. Keenan et al., 2016), between group members and non-group members (e.g. Waser, 1977), between kin and non-kin (e.g. Rendall et al., 1996), between neighbouring groups (Briseño-Jaramillo et al., 2015), and infer third-party rank relations (e.g. Borgeaud et al., 2013; Kitchen et al., 2005; Slocombe et al., 2010). Individually different calls have been described in many primate groups (e.g. blue monkeys: Butynski et al., 1992; Japanese macaques: Ceugniet & Izumi, 2004; spider monkeys Chapman & Weary, 1990; white-faced capuchin monkeys: Digweed et al., 2007; golden snub-nose monkeys: Fan et al., 2019; chacma baboons: Fischer et al., 2008; common marmosets: Jones et al., 1993; Miller et al., 2010; Wied’s black tufted-ear marmosets: Jorgensen & French, 2010; western gorillas: Salmi et al., 2014; cottom-top tamarins: Snowdon et al., 1983; gray-cheeked mangabeys: Waser, 1977). Yet, direct experimental evidence of whether individual variation in calls is perceived and used by group members is less frequent (but see e.g. in chimpanzees: Bauer & Michelle Philip, 1983; Kojima et al., 2003; vervet monkeys: Cheney & Seyfarth, 1980; Barbary macaques: Hammerschmidt & Fischer, 1998; common marmosets: Miller & Wren Thomas, 2012; rhesus monkeys: Rendall et al., 1996; cotton-top tamarins: Sproul et al., 2006; Weiss et al., 2001).

Spider monkeys from the genus *Ateles* inhabit semi-evergreen or tropical evergreen forest and range from southern Mexico to south eastern Brazil (Eisenberg, 1976). They are primarily frugivorous, feeding on fleshy fruits located in patches. They feed selectively at moderate to extreme heights in mature forests away from each other and contact calls are used to regroup after feeding (Chapman, 1988a, 1988b; Eisenberg, 1976). Spider monkeys live in groups of 15-40 members, but regularly split in subgroups (of 1-20 individuals) several times a day (Ramos-Fernández, 2005). Groups are highly fluid communities of multiple males and females. These groups show great variability in their party composition, party size and spatial cohesion amongst troop members (Boeving et al., 2017). During feeding cycles, spider monkeys tend to fractionate into unisex subgroups (Carpenter, 1935; Eisenberg, 1976). Within these subgroups during fractionation, males tend to be more independent than their female counterparts and during intra-troop conflicts offer mutual support (Eisenberg, 1976). Males tend to associate more with other males while females are more likely to associate with their more dependent young (Fedigan & Baxter, 1984).

When these spider monkeys are dispersed through wooded areas and are blocked from the sight of others, they emit loud calls also known as *whinnies*, a type of contact call, that are heard over long distances and are replied to with a similar whinny (Teixidor & Byrne, 1999). Whinnies can carry a distance of up to 300m which then subsequently elicit responses from the subgroup (Ramos-Fernández, 2005, 2008; Santorelli et al., 2013). These high frequency calls are used in different social context (feeding, moving resting), and are used to keep individuals in contact with each other together (Ordóñez-Gómez et al., 2018, 2019; Ramos-Fernández, 2005; Teixidor & Byrne, 1999). The ability to recognise each other by contact calls or *whinnies* can be extremely useful in relation to feeding and foraging as these whinnies can help locate monkeys that possess the best knowledge of whereabouts of resources (Chapman & Weary, 1990).

Given that contact calls (such as whinnies) play a role in intergroup social interactions, information about the identity of the caller is more likely than in other call types for which the caller identity is less important (e.g. alarm calls, or intergroup calls) (Santorelli et al., 2013). Several studies have shown that there is individual variation in the whinny calls of *Ateles geoffroyi* (Chapman & Weary, 1990; Ramos-Fernández, 2005; Santorelli et al., 2013; Teixidor & Byrne, 1997, 1999), suggesting the potential for individual recognition in the genus *Ateles*. However, even if calls differ between individuals, it does not necessarily mean the individual receivers perceive it or respond to it. Here we examined whether black-headed spider monkeys (*Ateles fusciceps*), a species whose vocal repertoire and social behaviour are very similar to *A. geoffroyi* (Eisenberg, 1976), could discriminate between familiar individuals using *whinny* calls, using a habituation-dishabituation paradigm.

## Methods

### Study system and site

Six black-headed spider monkeys (2 males, 4 females) from the Fota Wildlife Park, county Cork, Ireland, took part in the experiment (Table 1). Testing took place in their indoor enclosure, but subjects always had access to both their indoor and outdoor enclosure. For ethical reasons, individuals were never forcibly isolated from each other during the experiment.

**Table 1:**
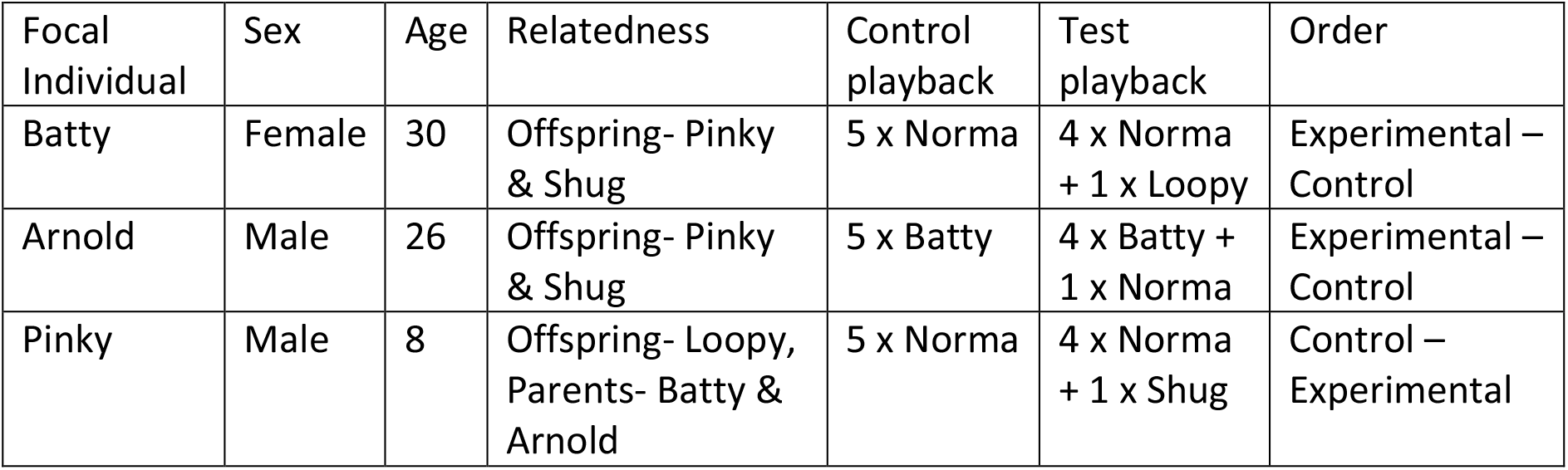
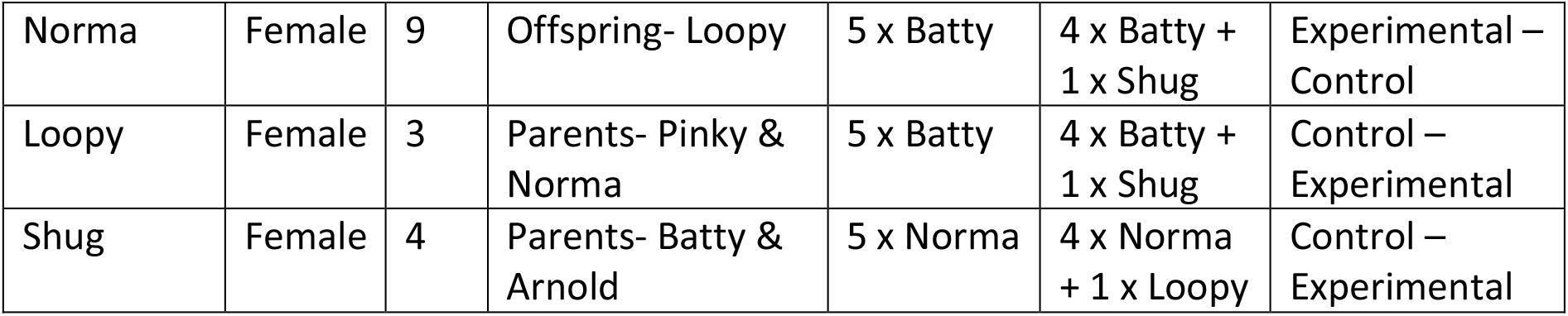
Age, sex and relatedness of the black-headed spider monkeys at Fota Wildlife Park, and the individuals used in the playback for each individual’s trials

| Focal Individual | Sex | Age | Relatedness | Control playback | Test playback | Order |
| --- | --- | --- | --- | --- | --- | --- |
| Batty | Female | 30 | Offspring- Pinky & Shug | 5 x Norma | 4 x Norma + 1 x Loopy | Experimental – Control |
| Arnold | Male | 26 | Offspring- Pinky & Shug | 5 x Batty | 4 x Batty + 1 x Norma | Experimental – Control |
| Pinky | Male | 8 | Offspring- Loopy, Parents- Batty & Arnold | 5 x Norma | 4 x Norma + 1 x Shug | Control – Experimental |
| Norma | Female | 9 | Offspring- Loopy | 5 x Batty | 4 x Batty +<br>1 x Shug | Experimental –<br>Control |
| Loopy | Female | 3 | Parents- Pinky &<br>Norma | 5 x Batty | 4 x Batty +<br>1 x Shug | Control –<br>Experimental |
| Shug | Female | 4 | Parents- Batty &<br>Arnold | 5 x Norma | 4 x Norma<br>+ 1 x Loopy | Control –<br>Experimental |

### Recording and extraction of whinnies

We first recorded *whinny* calls for each individual, using a Song Meter SM4 (Wildlife Acoustics). Calls were recorded between 10-11 am every day before feeding times, from the 31^st^ of October 2019 to the 11^th^ of November 2019, while the monkeys were freely allowed to roam in and out of the indoor enclosure. Calls were recorded from the indoor enclosure, where they could be easily observed, allowing us to control for distance to caller, and context of calling. Using the Audacity software, whinnies were identified from the recording based on sonograms. Studies on both *A. fusciceps* and *A. geoffroyi* were used to select whinnies based on the sonograms (*A. fusciceps*: Eisenberg, 1976; *A. geoffroyi*: Santorelli et al., 2013; Teixidor & Byrne, 1999), as whinnies of both species are very similar (Eisenberg, 1976), and vary between 2000 and 6000 Hz (Santorelli et al., 2013; Teixidor & Byrne, 1999), but sonograms of *A. geoffroyi* where of better quality in the literature.

### Habituation-dishabituation paradigm

The habituation-dishabituation paradigm is frequently used to study individual recognition, and is based on the ability to discriminate two stimuli (Cheney & Seyfarth, 1988; Johnston & Jernigan, 1994; Rendall et al., 1996). It requires habituation in response to repeated stimuli (e.g., different contact calls from the same individual), followed by a dishabituation phase in response to a different stimulus (e.g., one contact call from a different individual). As individuals repeatedly hear the same individual calling (using different calls), they should decrease their behavioural response (i.e., habituation phase). However, if individuals are able to discriminate between individuals based on their call, when a call from a different individual is played (i.e., dishabituation phase), individuals should increase their behavioural response (as it is no longer the same individual calling). If focal individuals do not increase their behavioural response, then this is interpreted as a failure to discriminate between two individuals based on the calls.

Due to the sex ratio of our sample size (two males : 4 females), only female whinnies were used to create playback. Playbacks consisted of 5 calls, separated by a 10 second interval. This was to allow individuals to react to the call before the next call was played. Control playbacks consisted of five different whinnies from the same individual, while experimental playbacks consisted of four different whinnies from the same individual and a fifth whinny from a different individual.

Once playbacks were created, the audio was normalised in Audacity, to set the peak amplitude of the playback. This set a constant amount of gain in the audio files to bring the amplitude to a normal level and did not affect sound quality of the recordings.

Playbacks were played before feeding time, which was just after 10 am everyday inside the enclosure. Each focal individual took part in two trials: one where they heard a control playback, and one where they heard an experimental playback (Table 1). The order in which trials took place was randomly assigned (Table 1). The focal individual was never tested more than once each day to avoid carry-over habituation effects on the same day. Moreover, although we conducted trials when few individuals were in the indoor enclosure, since trials could not be conducted in complete isolation, not more than three trials a day were performed on the group to avoid habituation.

When the focal individual was identified, the corresponding playback (either control or experimental) was played directly from Audacity from a laptop to an EasyAcc speaker connected via Bluetooth, located just outside of the enclosure. The volume was set to a consistent level for each trial. The noise level was selected to reflect the natural level of a whinny. Each trial was filmed with a Panasonic HC-W580 camera. Following Teixidor & Byrne (1997), we recorded the following binary variables (1) whether the focal individual looked at the speaker during the call or during the 10 second interval after the call, (2) whether they approached the speaker, (3) if they vocalised in response of the playback, (4) if there were any physical interactions with other individuals during the playback, and (5) whether they displayed aggressive behaviour (e.g. rattling the cage, banging against the cage or initiating physical aggressive contact with another individual). We also recorded the amount of time spent looking at the speaker.

### Statistical analysis

Most binary variables showed little variation (individuals did not look at the speaker in 6/60 trials, interacted with other individuals on 2/60 trials, vocalized on 3/60 trials and showed aggression on 4/60 trials). Analyses were therefore conducted only on time spent looking at the speaker and whether individuals approached the speaker or not.

All analyses were conducted in R (R Core Team, 2022). To investigate whether the interaction between playback type (experimental or control) and call number within the playback (1 to 5) had an effect on our behavioural measures, we ran generalised linear mixed models using the *glmer* function of the *lme4* package (Bates et al., 2015). We included the treatment (control or experimental playback), call number within the playback (1-5), as well as the interaction between call number and treatment, as fixed effects. We also controlled for age and sex of the individual, as well as the order of playback (first vs second session), by including them as fixed effects. Finally, we also included focal individual as a random effect. We used a Poisson error structure for examining the time spent looking at the speaker, and a binomial error structure for examining whether individuals approached the speaker or not. We then ran a posthoc test to examine the effect of the interaction on our behavioural response, using the *emmeans* package (Lenth, 2019). Model assumptions were checked using the *DHARMa* package (Hartig, 2020). We used the *cat_plot* function from the *interactions* package and the *effect_plot* function from the *jtools* package to plot model effects (Long, 2019, 2022).

### Data availability

Data and code are available on OSF: https://osf.io/z698r/?view_only=95c4f0365a4549999d01675664250f2e

### Ethics

We performed the experiment in accordance with the Association for the Study of Animal Behaviour guidelines for the Treatment of Animals in Behavioral Research and Teaching. The study was approved by Fota Wildlife Park’s ethical committee.

## Results

Individuals spent on average 4.2 seconds (range: 0-10 seconds) looking at the speaker during the trials, and approached the speaker on 13/60 trials.

There was a significant effect of the interaction between playback type and the call number within the playback (Table 2, Figure 1). During the first, second, third and fourth call, there was no significant difference in time spent looking at the speaker between the control and experimental playback (Table 2, 3, Figure 1). However, during the fifth call, there was a significant difference between the two playback type: individuals looked more at the speaker during the fifth call of the experimental playback than during the fifth call of the control playback (Table 3, Figure 1).

**Table 2:** Generalised linear mixed models, comparing the time spent looking for each playback type (control vs experimental), call number within the playback (1-5), age, sex (female vs male), order of playback, and the interaction between playback type and trial number. Individual id was included as a random effect. ^1^ baseline = control; ^2^ baseline = call 1; ^3^ baseline = female; ^4^ baseline = 1

| Variable | Estimate (Std. Error) | 95% CI | p-value |
| --- | --- | --- | --- |
| **Intercept** | **2.00 (0.200)** | **1.60; 2.39** | **<0.001** |
| Playback: Experimental<sup>1</sup> | -0.241 (0.220) | -0.673; 0.187 | 0.273 |
| Call 2<sup>2</sup> | -0.213 (0.218) | -0.643; 0.212 | 0.329 |
| **Call 3<sup>2</sup>** | **-0.672 (0.251)** | **-1.18; -0.195** | **0.007** |
| **Call 4<sup>2</sup>** | **-1.29 (0.313)** | **-1.94; -0.702** | **<0.001** |
| **Call 5<sup>2</sup>** | **-2.06 (0.433)** | **-3.01; -1.29** | **<0.001** |
| Age | -0.002 (0.008) | -0.022; 0.016 | 0.808 |
| Sex: Male<sup>3</sup> | 0.287 (0.179) | -0.128; 0.711 | 0.110 |
| Order: 2<sup>4</sup> | -0.049 (0.126) | -0.296; 0.197 | 0.699 |
| Playback x Call (Experimental x Call 2) | 0.036 (0.327) | -0.606; 0.676 | 0.913 |
| Playback x Trial (Experimental x Call 3) | -0.300 (0.401) | -1.10; 0.479 | 0.455 |
| Playback x Trial (Experimental x Call 4) | 0.159 (0.456) | -0.745; 1.06 | 0.727 |
| **Playback x Trial (Experimental x Call 5)** | **1.85 (0.498)** | **0.947; 2.90** | **<0.001** |

**Table 3:** Posthoc test for the linear model results for time spent looking between the control and experimental playback for each call within the playback

| Call Number | Estimate (std. error) | 95% C.I. | p-value |
| --- | --- | --- | --- |
| 1 | 0.241 (0.220) | -0.190; 0.672 | 0.273 |
| 2 | 0.205 (0.242) | -0.269; 0.679 | 0.396 |
| 3 | 0.541 (0.336) | -0.118; 1.20 | 0.108 |
| 4 | 0.082 (0.400) | -0.702; 0.866 | 0.839 |
| 5 | **-1.61 (0.447)** | **-2.48; -0.732** | **<0.001** |

**Figure 1:**
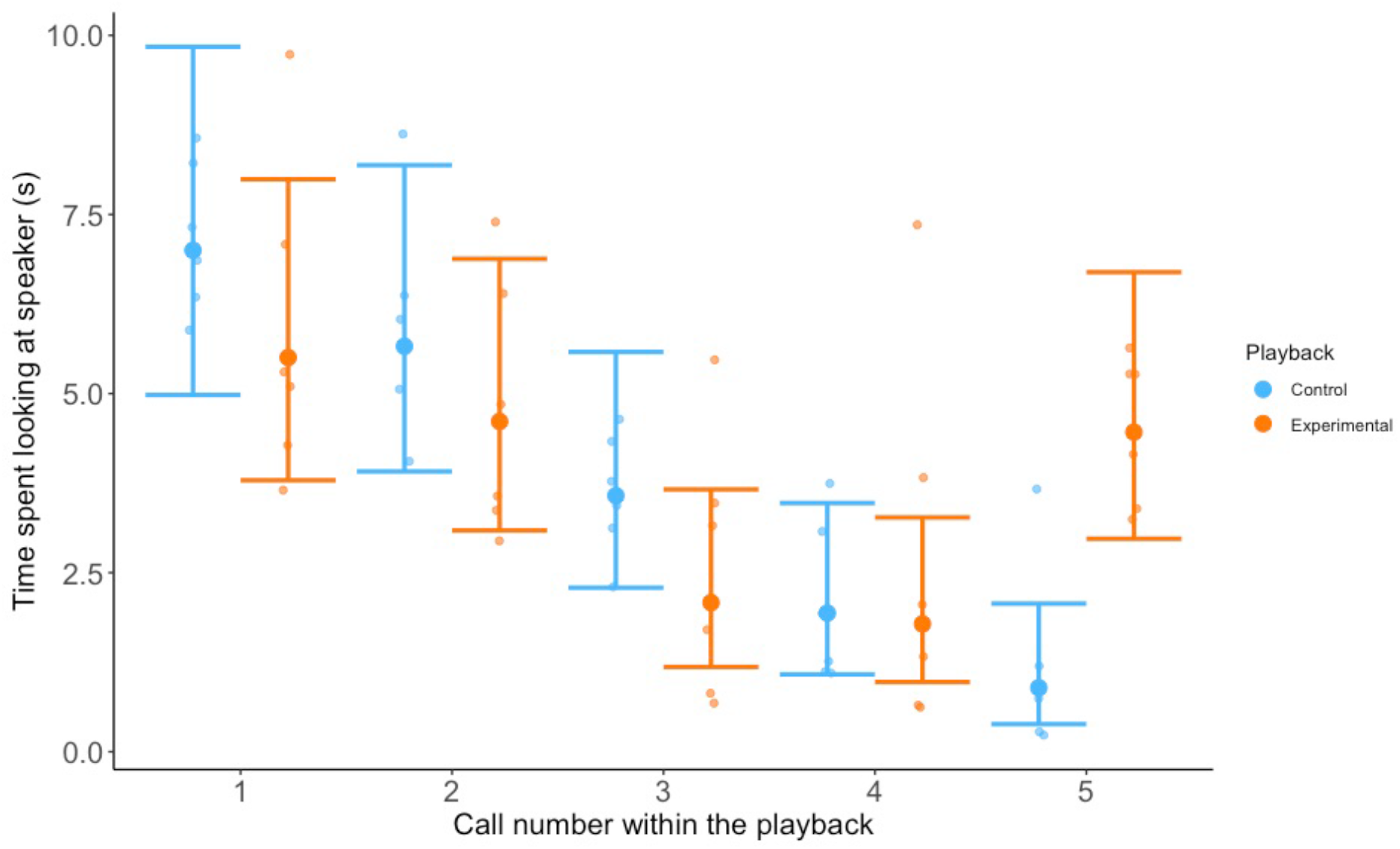
Partial residual plots showing time spent looking at the speaker across the five calls within the playback, for the control and experimental playbacks Error bars are 95% confidence intervals based on the linear mixed models in Table 2.

Males tended to look at the speaker for longer compared to females, but this difference was not statistically significant (Table 2). There was no evidence of a significant effect of age of the focal individual and order of the playback, on the time spent looking at the speaker (Table 2).

Males were more likely to approach the speaker than females (Table 4). There was no evidence of a significant effect of the interaction between playback type and call number, age of the focal individual, and order of the playback on whether individuals approached the speaker or not (Table 4).

**Table 4:** Generalised linear mixed models, comparing whether individuals approached the speaker for each playback type (control vs experimental), call number within the playback (1-5), age, sex (female vs male), order of playback, and the interaction between playback type and trial number. Individual id was included as a random effect. ^1^ baseline = control; ^2^ baseline = call 1; ^3^ baseline = female

| Variable | Estimate (Std. Error) | 95% CI | p-value |
| --- | --- | --- | --- |
| Intercept | 0.183 (1.19) | -2.16; 2.52 | 0.878 |
| Playback:<br>Experimental<sup>1</sup> | -0.828 (1.40) | -3.57; 1.92 | 0.554 |
| Call 2<sup>2</sup> | -0.981 (1.41) | -3.74; 1.77 | 0.485 |
| Call 3<sup>2</sup> | -0.982 (1.41) | -3.74; 1.77 | 0.485 |
| Call 4<sup>2</sup> | -22.0 (1.66e4) | -3.26e4; 3.25e4 | 0.999 |
| Call 5<sup>2</sup> | -22.0 (1.73e4) | -3.36e4; 3.36e4 | 0.999 |
| Age | -0.045 (0.051) | -0.145; 0.056 | 0.384 |
| **Sex: Male<sup>3</sup>** | **2.47 (1.06)** | **0.398; 4.53** | **0.019** |
| Order | -0.909 (0.814) | -2.50; 0.687 | 0.264 |
| Playback x Call<br>(Experimental x Call<br>2) | -0.205 (2.11) | -4.33; 3.92 | 0.922 |
| Playback x Trial<br>(Experimental x Call<br>3) | -0.205 (2.11) | -4.33; 3.92 | 0.922 |
| Playback x Trial<br>(Experimental x Call<br>4) | 0.419 (2.76e4) | -5.40e4; 5.40e4 | 1.00 |
| Playback x Trial<br>(Experimental x Call<br>5) | 22.0 (1.71e4) | -3.36e4; 3.36e4 | 0.999 |

## Discussion

Individuals spent less time looking at the speaker, that is, they habituated to the caller, as the number of calls from the same individuals increased during the playback. However, they looked significantly more at the speaker when the fifth call was that of another individual (experimental playback), compared to when the fifth call was that of the same individual (control playback). This supports the hypothesis that spider monkeys can identify individuals through contact calls and agrees with the assumption made in wild studies that whinnies are used to stay in contact with known group members (Ordóñez-Gómez et al., 2018, 2019; Ramos-Fernández, 2005; Teixidor & Byrne, 1999). Fission-fusion is a major aspect of sociality in *Ateles* and this may be the main driving force behind the ability to recognise individuals through vocal cues (Aureli & Schaffner, 2008), helping individuals find group members foraging outside their own subgroup, help them decide which subgroup to join, or know how many individuals are depleting a food patch (Chapman & Weary, 1990; Teixidor & Byrne, 1999).

We found little behavioural response other than looking towards the speaker, which is also in line with previous research in spider monkeys (Teixidor & Byrne, 1997, 1999). Whinny recognition does not necessarily require a strong change in behaviour, especially if individuals are in close proximity. Studies also found that different responses are elicited depending on the context in which the whinnies were used (Teixidor & Byrne, 1999). It is likely that the small group size our study was based on did not provide sufficient variation in the sort of context in which stronger responses would be elicited e.g. separation of subgroups for prolonged periods of time or rivalry between males.

Males tended to look at the speaker for longer, compared to females (although the difference was not statistically significant) and males were more likely than females to approach the speaker. Males are the more gregarious sex and females are more likely to be found alone (Aureli & Schaffner, 2008). Females travel in smaller groups compared to males, except when food is abundant, and groom each other less (Aureli & Schaffner, 2008). Males in comparison might therefore respond more strongly to whinnies in order to join a group, which might explain why they were more likely to approach the speaker, regardless of the identity of the individual calling.

Previous recording and playback experiments showed that spider monkeys respond differently to stranger and familiar whinnies (Teixidor & Byrne, 1997) and to whinny calls emitted in different contexts (Teixidor & Byrne, 1999), but they do respond consistently to the same call (Masataka, 1986). Although there is variation in whinny calls between independent social groups (Santorelli et al., 2013), and between context (Teixidor & Byrne, 1999), most of the variation in the spider monkey calls across groups is at the individual level (Chapman & Weary, 1990; Ramos-Fernández, 2005; Santorelli et al., 2013; Teixidor & Byrne, 1997, 1999). Here we have shown that individuals can use this individual variation in the whinny call to discriminate between individual callers. However, not all individual responded similarly, which could be explained by differences in the identity of the caller and responder (Ramos-Fernández, 2005; Teixidor & Byrne, 1999), or the relationship between the caller and responder (Masataka, 1986; Ramos-Fernández, 2005). Given that spider monkeys can discriminate between individual whinnies, the next step would be to know in which contexts they use individual identity of caller to make decisions.

## Acknowledgements

We would like to thank the members of Fota Wildlife Park staff, particularly Sean McKeown, for allowing access to animals.

## AUTHOR CONTRIBUTIONS

Conceptualization: C.A.T, with input from D.N and J.L.Q. Data curation: D.N and C.A.T. Formal analysis: D.N and C.A.T. Investigation: D.N with input from T.P. Methodology: D.N and C.A.T. Project administration: D.N. Supervision: C.A.T. Writing – original draft: D.N and C.A.T. Writing – review and editing: D.N, T.P, J.L.Q, C.A.T.

